# Modeling time-varying phytoplankton subsidy reveals at-risk species in a Chilean intertidal ecosystem

**DOI:** 10.1101/2023.07.27.550852

**Authors:** Casey S. Duckwall, John L. Largier, Evie A. Wieters, Fernanda S. Valdovinos

## Abstract

The allometric trophic network (ATN) framework for modeling population dynamics has provided numerous insights into ecosystem functioning in recent years. Herein we extend ATN modeling of the intertidal ecosystem off the central coast of Chile to include empirical data on pelagic chlorophyll-a concentration. This intertidal community requires subsidy of primary productivity to support its rich ecosystem. Previous work models this subsidy using a constant rate of phytoplankton input to the system. However, data shows pelagic subsidies exhibit short-term, highly variable, pulse-like behavior. Incorporating this variable input into ATN modeling is the primary contribution of this work and provides several new insights into this ecosystem’s response to pulses of offshore phytoplankton, including: (1) closely related sea snails show differential responses to pulses of phytoplankton that are explained by underlying network structure; (2) increasing the rate of pelagic-intertidal mixing increases fluctuations in species’ biomasses that may increase the risk of local extirpation; (3) predators are the most sensitive species to phytoplankton biomass fluctuations, putting these species at greater risk of extirpation than others. Finally, our work provides a straightforward way to incorporate empirical, time-series data into the ATN framework that will expand this powerful methodology to new applications.

## Introduction

Oceanic subsidies of phytoplankton are essential to the persistence of near-shore intertidal communities [1]. In upwelling systems, increased phytoplankton subsidy increases recruitment, growth, and reproduction of invertebrate populations [2–5], which increases the secondary productivity transferred to higher trophic levels in intertidal food webs [6–8]. These subsidies, however, are highly variable due to complex dynamics of wind-driven upwelling that control phytoplankton blooms [9, 10], which are modulated by seasons, interannual variability, and local geography, as well as by wave-driven circulation that contributes to the delivery of phytoplankton to intertidal habitats [11–13]. This complex variability in phytoplankton subsidies forces non-autonomous dynamics (i.e., explicitly dependent on time) into the productivity of intertidal food webs, which challenges the autonomous approach traditionally used to model food web dynamics (i.e., dependent only on fixed parameters representing the local trophic and demographic rates). These non-autonomous factors will most likely cause intertidal food webs to be dominated by transient dynamics [14], as opposed to equilibrium dynamics, by repeatedly pushing the system away from any trajectory approaching equilibrium. Understanding the effects of these non-autonomous dynamics on ecological systems is an important new frontier in community ecology given the dramatic environmental changes currently altering ecosystem dynamics worldwide.

The Humboldt Current System supports a nutrient-rich and high-diversity ecosystem off the coast of Chile and Peru [15]. These waters are among the most productive marine ecosystems in the world [16, 17] and exhibit dramatic annual fluctuations in phytoplankton abundance due to several, often interacting forces [18]. The high productivity of this system is supported by wind-driven upwelling, a bulk transport process that forces surface water offshore to be replaced with deep, nutrient-rich water [9, 19–21]. Upwelling favorable winds exhibit spatiotemporal variability, driving synoptic (shorter-term fluctuations lasting only for days or weeks) and seasonal cycles in upwelling, nutrient concentration, and oceanic phytoplankton density [9, 22]. Additionally, global oceanographic and meteorological phenomena, such as the El Niño-Southern Oscillation and the Pacific Decadal Oscillation, alter productivity by introducing environmental variability on time scales of 4-7 and 20-30 years, respectively [23, 24]. Finally, wind-driven upwelling is being modified by global climate change [17, 25, 26], resulting in significant changes in coastal phytoplankton concentrations [27] – further, pelagic-intertidal coupling can also be expected to change with changes in winds, waves and stratification that control exchange between the coastal ocean and shoreline habitats. This complex array of interacting processes makes the variability of oceanic phytoplankton subsidizing intertidal communities extremely difficult to predict or model from first principles. Therefore, we propose the use of empirical oceanic phytoplankton time series data to represent the variability of the phytoplankton subsidizing intertidal food webs. Specifically, our present contribution uses a time series of oceanic phytoplankton (measured as chlorophyll-a) from the central coast of Chile to evaluate the effects of such variability on the dynamics of a rocky intertidal food web.

The bioenergetic model [28] expanded to large food webs by the Allometric Trophic Network (ATN) model [29] has been successful in capturing the dynamic processes of aquatic food webs, from predicting interaction strengths of rocky-intertidal food webs [30] to modeling food web dynamics of lakes [31]. This model uses biomass (energy) as a currency and has been greatly influential toward understanding mechanisms behind community function and stability [32, 33] and for predicting consequences of environmental changes and human exploitation on diversity [6, 34–38]. A main reason for its success is that the ATN model leverages several trophic and metabolic processes that scale with the body mass of aquatic organisms [39–42], allowing researchers to use allometric scaling to parameterize the model for any system where body masses of the interacting species are known. This model, however, only accounts for the autonomous dynamics that emerge from allometrically scaled demographic and interaction rates. Previous work shows how the dynamics of the ATN model applied to the rocky intertidal system we study here are affected by including phytoplankton subsidy at a constant rate (fixed throughout the entire simulation), and subsequently increasing or decreasing this rate based on expectations from different climate change scenarios [6]. Here, we advance the field by modeling these subsidies more realistically by incorporating their high temporal variability, using time-series of chlorophyll-a to incorporate the empirical variability of oceanic phytoplankton into the subsidies received by the rocky intertidal food web.

Our specific objectives are to evaluate how the rocky intertidal food web of the central coast of Chile responds to (1) the variability of offshore phytoplankton abundance and (2) the rate of pelagic-intertidal mixing. These two factors combine to determine the flux of particulate food from the ocean to intertidal habitats. We found that while this pelagic subsidy is essential for fueling the intertidal food web, the high level of temporal variability (enhanced by rapid pelagic-nearshore exchange) increases biomass fluctuations putting species at increased risk of extirpation, and that variation in abundance due to phytoplankton subsidy is most pronounced in predators relative to other types of species.

## Results

Figure 1 outlines the study setup. Open-water chlorophyll-a measurements were obtained for the research site near Las Cruces, Chile (Fig. 1A). These empirical values were scaled to model units, and a spline curve was fit to the data (Fig. 1B) to create a continuous curve of offshore phytoplankton biomass. A network representation of the intertidal ecosystem is provided in Fig. 1C with nodes representing the various intertidal species and links representing their associated trophic interactions. The height of a species’ node corresponds to its trophic level. Variable pelagic phytoplankton (OP) drives this food web network by subsidizing phytoplankton in the intertidal habitat (FP). The rate constant, k_mixing_, controls the rate at which pelagic phytoplankton is mixed into the food web network. k_mixing_ is assigned values of 0.1, 1.0, and 10hr^−1^ as denoted. For ease of illustration, we will focus on results from 2003; however, trends are consistent across all years investigated. Results from other years are included in the supplementary materials and referenced where relevant. Summary information for the empirical dataset is provided in the supplementary materials (Supplementary Tables S1, S2).

**Figure 1 –.**
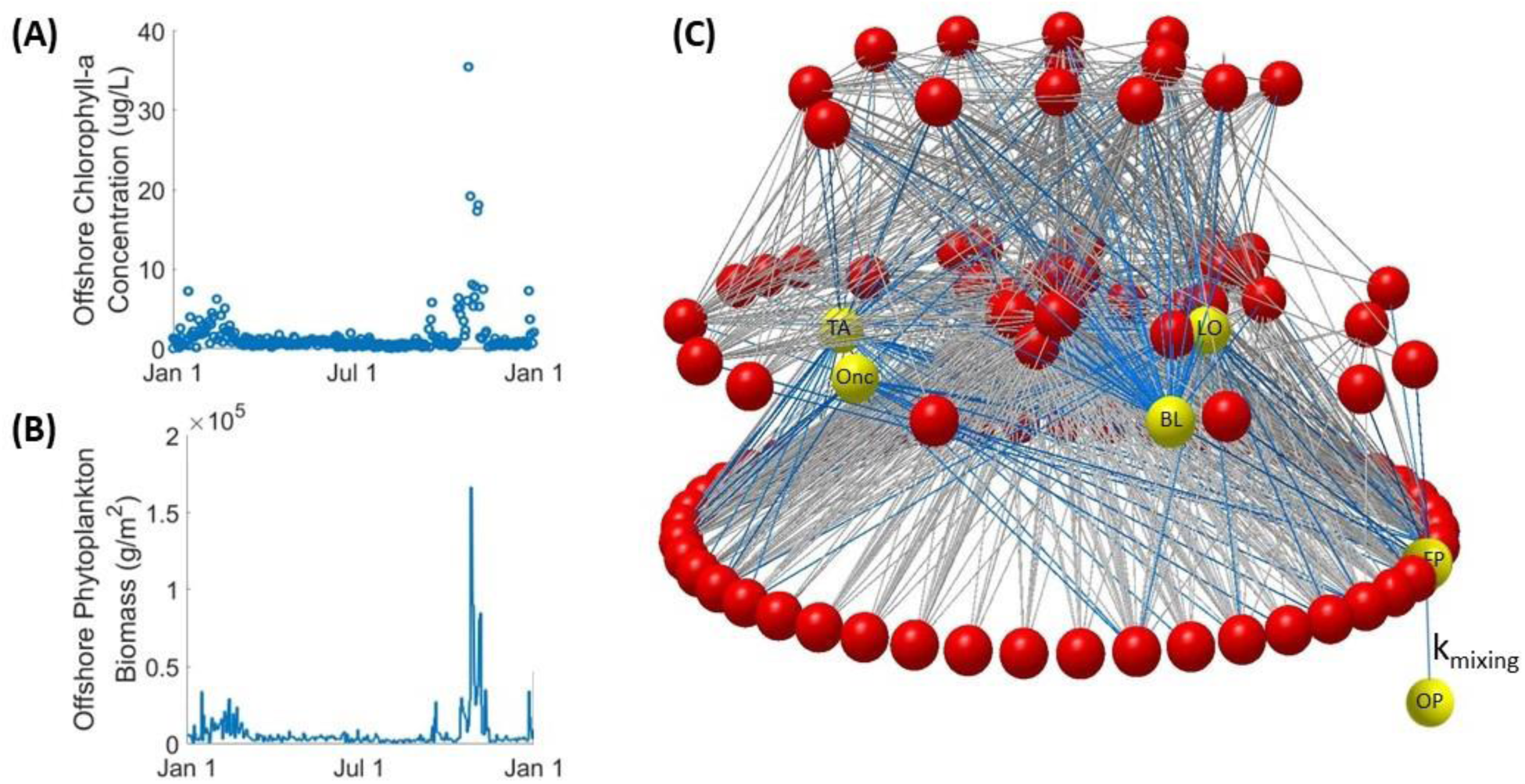
(A) Empirical timeseries dataset of offshore chlorophyll-a measurements in 2003. (B) Continuous spline fit to scaled empirical dataset. (C) Graphical representation of the Las Cruces food web network. Nodes are vertically arranged by using prey-averaged trophic level. Important nodes are labeled as follows: OP – offshore phytoplankton, FP – food web phytoplankton, BL – the barnacle, *B. laevis*, Onc – the sea snail, *Onchidella sp.*, TA – the sea snail, *T. atra*, LO – the sea snail, *L. orbignyi*.

### Species’ biomasses respond differently to phytoplankton variability depending on their food web connections

We began our investigation into the effects of pulses of high pelagic phytoplankton on intertidal population dynamics using an intermediate rate of pelagic-intertidal mixing, k_mixing_ = 1.0hr^−1^.

Pulses of high pelagic phytoplankton that occurred in late 2003 (labeled OP in Fig. 2A) are characteristic of seasonal upwelling in the central Chilean coast [22]. We found that the abundances of all filter feeder species were strongly and positively influenced by increased phytoplankton subsidy (Supplementary Fig. S2). Our model assumes that phytoplankton is the only resource available to filter feeders. Consequently, the biomasses of filter feeder species are tightly coupled with the abundance of intertidal phytoplankton. Figure 2B illustrates this result for the filter-feeding barnacle, *B. laevis*, an important food source for omnivore sea snails.

**Figure 2 –.**
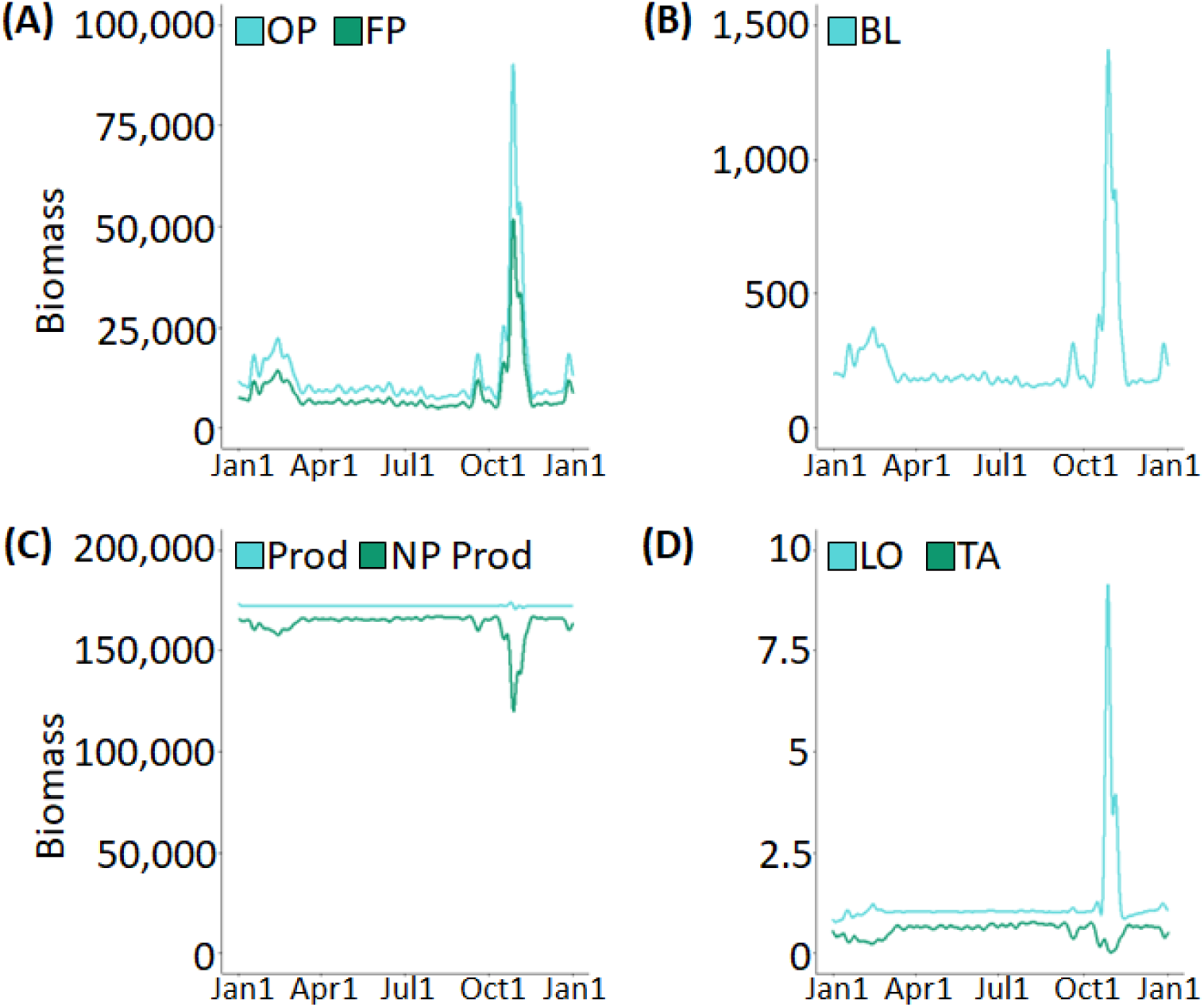
**Biomass curves simulated using empirical data of 2003 for** (A) offshore phytoplankton (OP) and food web phytoplankton (FP), (B) the barnacle, *B. laevis* (BL), (C) summed biomass of all producer species (Prod) and all non-planktonic producer species (NP Prod), and (D) the omnivorous sea snail, *L. orbignyi* (LO), and the herbivorous sea snail, *T. atra* (TA). Biomasses are shown in g/m^2^. These results use an intermediate pelagic-intertidal mixing rate, k_mixing_ = 1.0hr^−1^.

The total biomass of all producer species (including phytoplankton) remained relatively constant across the year (labeled Prod in Fig. 2C), but the total biomass of all non-planktonic producer species (i.e., algae) was negatively influenced by increased phytoplankton subsidy (labeled NP Prod in Fig. 2C). These two results are explained by the community-level carrying capacity that is shared by all producer species of the model (see Eq. 3). This carrying capacity limits the total biomass of producers so if some producer species increase their biomass others will decrease theirs due to competition for space or other resources.

Responses among consumer species were more varied. For example, the different sea snail species exhibited different responses to the Oct.-Nov. 2003 elevation in phytoplankton based on their feeding preference as either omnivore or herbivore (Supplementary Fig S3). On the one hand, omnivorous sea snails like *L. orbignyi* (LO in Fig 2D) directly consume filter-feeding barnacles, placing them two nodes above food web phytoplankton. Consequently, omnivorous sea snails were subject to variations in biomass similar to those observed in phytoplankton and their filter feeder prey. On the other hand, herbivorous sea snails like *T. atra* (TA in Fig 2D) consume algae and kelp but not barnacles nor any other consumers of phytoplankton. Consequently, increased phytoplankton affected herbivorous sea snails only indirectly by increasing the abundance of their predators and suppressing the growth of their resources. Through this example, we observe that offshore phytoplankton alters both top-down and bottom-up forces differentially even for related (within class) species and that these differences can be made evident through inspection of their network connections.

### Increased pelagic-intertidal mixing increases biomass fluctuations putting species at increased risk of extirpation

We simulated the effects of varying the pelagic-intertidal mixing rate on the rocky intertidal ecosystem by assigning values of 0.1, 1.0, and 10hr^−1^ to k_mixing_ (Fig. 3A,B). Although increasing values of k_mixing_ increased the concentration of phytoplankton in the rocky intertidal, our simulations showed only a small increase in total primary productivity due to the community-level carrying capacity shared by producer species (Fig. 3C). However, total consumer biomass – and biomass of filter-feeding invertebrates in particular – increased with k_mixing_, highlighting the importance of phytoplankton in fueling the intertidal ecosystem across trophic levels.

**Figure 3 –.**
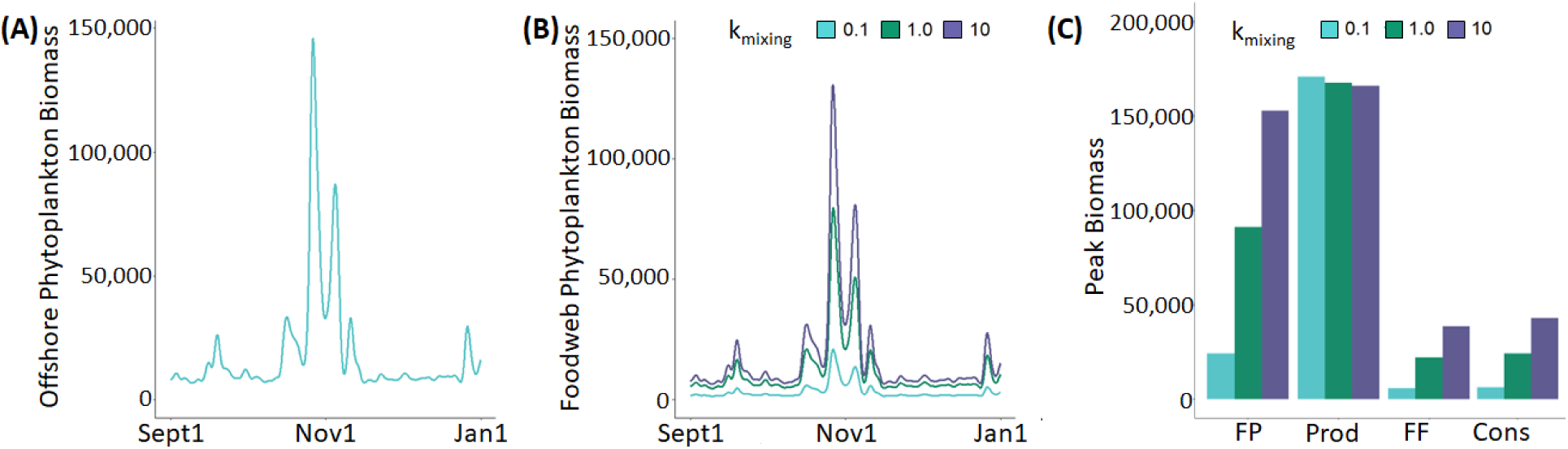
(A) Measured biomass of offshore phytoplankton from Sept 2003 to Jan 2004. (B) Simulated biomass of food web phytoplankton under three values of k_mixing_. (C) Peak simulated biomass from Sept 2003 to Jan 2004 for four trophic categories and three values of k_mixing_. FP – Food web phytoplankton biomass, Prod – Summed biomass of all 46 producer species, FF – Summed biomass of all 15 filter feeder species, Cons – Summed biomass of all 60 consumer species.

The pelagic-intertidal mixing rate exhibits profound influence on the persistence of certain species, as highlighted by the response of the sea snail *Onchidella* to the peak in offshore phytoplankton during Oct.-Nov. 2003 (Fig. 4A). Using an intermediate mixing rate (k_mixing_ = 1.0hr^−1^), our model predicts that the biomass of *Onchidella* would have reached a critical threshold in late October 2003 (Fig. 4B – green) caused by an increase in offshore phytoplankton that produced a moderate reduction in its resources (microalgae and kelp, Fig. 4C – green) and a large increase in its predator biomass (Fig. 4D – green). The ATN model considers a species at risk of local extirpation when its calculated biomass drops below a critical threshold – in our case 10^−6^ g/m^2^ [6, 33] – because such extremely low biomass levels do not recover without an external input of biomass of such species.

**Figure 4 –.**
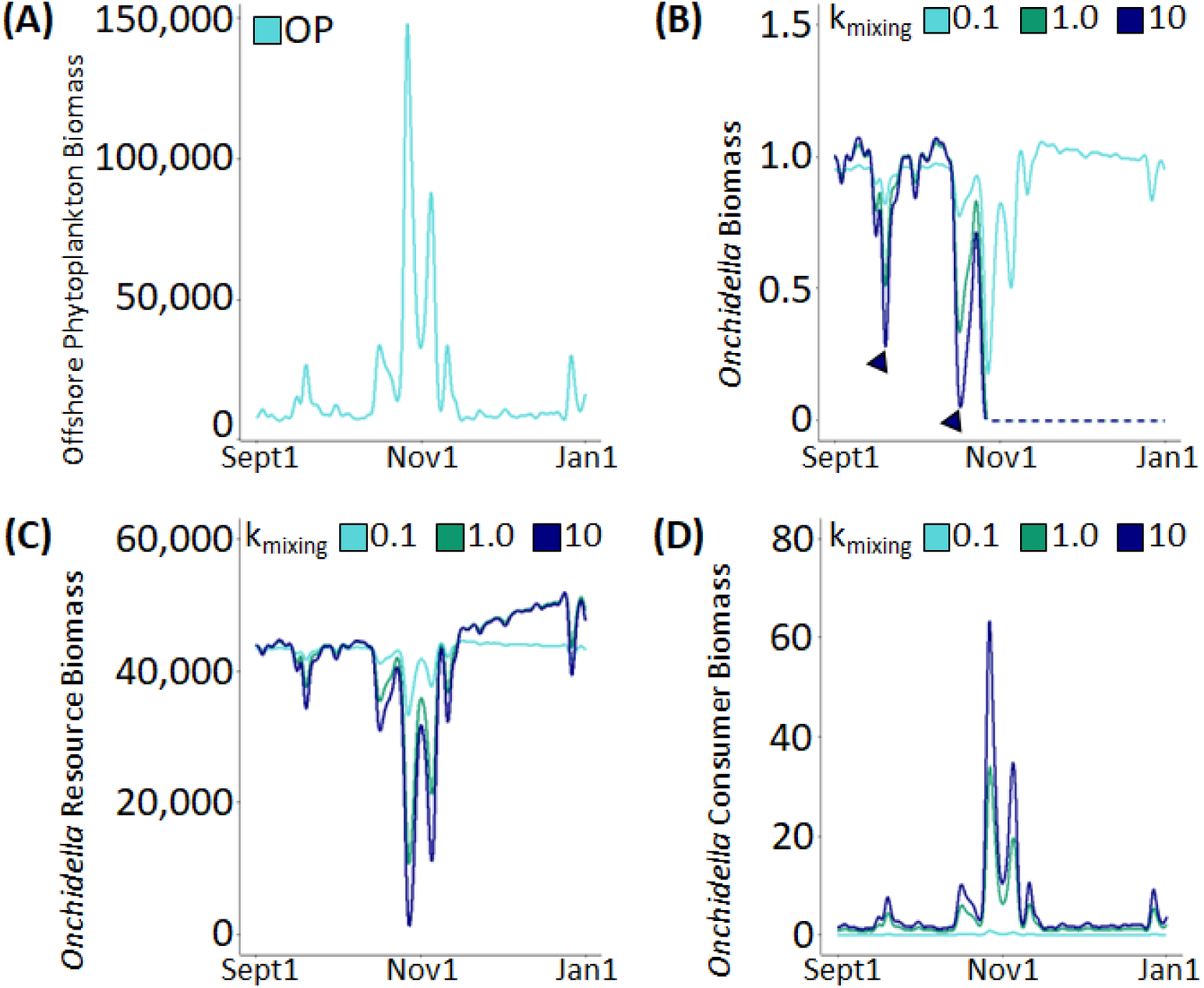
Results shown span Sept 2003 to Jan 2004 and show three values of k_mixing_. Biomass given in g/m^2^. (A) Measured biomass of offshore phytoplankton. (B) Simulated biomass of sea snail *Onchidella*. (C) Sum of simulated biomasses for the 14 species *Onchidella* eats. (D) Simulated biomass of the crab *Acanthocyclus gayi*, *Onchidella*’s sole known predator in Las Cruces.

Decreasing the mixing rate to 0.1hr^−1^ results in slower exchange between intertidal habitat and offshore waters and rescues *Onchidella* from extirpation (Fig. 4B – teal) because the biomasses of its resources (Fig. 4C – teal) and predator (Fig. 4D – teal) are significantly less affected by offshore phytoplankton. Conversely, increasing the mixing rate to 10hr^−1^ results in almost immediate exchange with offshore waters and exacerbates the extirpation that was observed for intermediate mixing with both a stronger reduction in *Onchidella*’s resources (Fig. 4C – navy) and a more dramatic increase in its predator biomass (Fig. 4D – navy). The biomass of *Onchidella* itself was much more variable under higher mixing rates with two additional incidents when *Onchidella* was at critical risk of extirpation (Fig 4B – navy, triangles).

In our simulations, the survival of *Onchidella* was influenced by two factors: the peak abundance of offshore phytoplankton and the choice of pelagic-intertidal mixing rate. Our simulations indicate (1) that *Onchidella* can survive during periods with relatively modest peaks in offshore phytoplankton at all rates of pelagic-intertidal mixing, and (2) that *Onchidella* can survive scenarios with low pelagic-intertidal mixing rates regardless of the intensity of fluctuations in offshore phytoplankton abundance (Supplementary Fig. S4, Table 1). However, during years with intermediate peaks in offshore phytoplankton, *Onchidella* faced extirpation when subjected to high rates of pelagic-intertidal mixing (k_mixing_ = 10hr^−1^). Further, during years with very high peaks in offshore phytoplankton abundance, *Onchidella* experienced extirpation at both high (k_mixing_ = 10hr^−1^) and intermediate (k_mixing_ = 1.0hr^−1^) levels of pelagic-intertidal mixing. We observe that *Onchidella’s* survival is modulated in a dose-dependent manner by peak abundance of offshore phytoplankton and that an appropriate combination of peak offshore phytoplankton abundance and pelagic-intertidal mixing rate is necessary for its survival.

**Table 1 –.** Results list peak offshore phytoplankton abundance and simulated extirpation status of *Onchidella* under each pelagic-intertidal mixing scenario investigated. Cells with an ‘e’ signify simulations in which *Onchidella* experienced extirpation, and empty cells signify simulations where *Onchidella* survived.

| <u>Year</u> | <u>Peak OP Abundance</u> | <u>Status Under</u> |  |  |
| --- | --- | --- | --- | --- |
|  |  | <u>$k_{\text{mixing}} = 10$</u> | <u>$k_{\text{mixing}} = 1$</u> | <u>$k_{\text{mixing}} = 0.1$</u> |
| 1999 | 297,260 | e | e |  |
| 2000 | 191,120 | e | e |  |
| 2001 | 47,860 | e |  |  |
| 2002 | 131,400 | e | e |  |
| 2003 | 172,440 | e | e |  |
| 2004 | 50,338 | e |  |  |
| 2005 | 64,821 | e |  |  |
| 2006 | 74,076 | e | e |  |
| 2007 | 179,560 | e | e |  |
| 2008 | 61,371 | e |  |  |
| 2009 | 27,577 |  |  |  |
| 2010 | 17,871 |  |  |  |

### Variation from phytoplankton subsidy is most pronounced in consumers and is exacerbated by higher pelagic-intertidal mixing rates

Within-year variations in species biomasses caused by fluctuations in offshore phytoplankton concentration differed across species guilds. To assess the variability across the network, each species was assigned to one of five guilds: filter feeders, producers, herbivores, omnivores, and carnivores. Variability within each guild was evaluated using both a species-specific normalized coefficient of variation (Fig 5) and a guild-summed normalized coefficient of variation (Supplementary Table S4).

**Figure 5 –.**
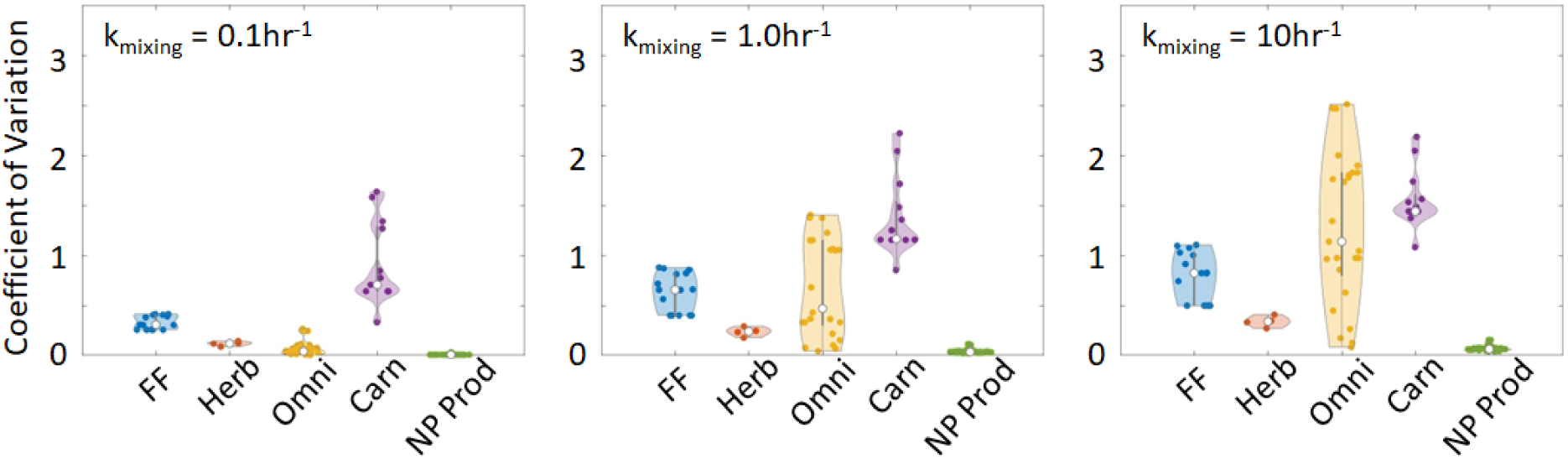
**Violin plots showing the coefficients of variation for each species normalized to that of offshore phytoplankton.** Results shown for 2003 and each value of k_mixing_ investigated. Species were grouped into one of five guilds: filter feeders (FF), herbivores (Herb), omnivores (Omni), carnivores (Carn), and non-planktonic producers (NP Prod).

For each given rate of pelagic-intertidal mixing, the non-planktonic producer guild exhibited the lowest normalized coefficient of variation relative to offshore phytoplankton. The high biomass of algae and kelp causes the variation-to-mean ratio in non-phytoplankton producers to be small relative to that of phytoplankton. Herbivore species also exhibited relatively low normalized coefficients of variation, explained by their tight trophic relationship with non-planktonic producer species.

Filter feeders, omnivores, and carnivores exhibited differential patterns of variability under different scenarios of pelagic-intertidal mixing. As the pelagic-intertidal mixing rate parameter was raised, variability in the biomasses of filter feeders increased. This can be attributed to two factors: (1) the strong trophic connection between filter feeders and the abundance of phytoplankton in the food web and (2) our previous result showing that higher values of pelagic-intertidal mixing rate enhance the availability of phytoplankton as a food source for filter feeders. Together, these two factors will push variability in filter feeder abundances closer to that of offshore phytoplankton. Variability arising from fluctuations in offshore phytoplankton abundance is more ambiguous in omnivore species. Omnivores exhibit extremely low relative variability under low pelagic-intertidal mixing rates, with relative variability increasing dramatically with mixing rate. Variability in carnivore species’ biomasses, however, shows elevated relative variability even in scenarios with low pelagic-intertidal mixing rate. Further, carnivores show a clear trend of increased variability as a result of increased pelagic-intertidal mixing rate. Carnivores tend to have low total biomasses and incorporate several different sources of variability due to their position near the top of the food web, causing their biomass variability to be large. These observations are consistent with previous studies showing that predators amplify biomass fluctuations and exhibit high sensitivity to network perturbations [43].

## Discussion

The rich and diverse ecosystems of the rocky shore would not be possible without the subsidy to their primary productivity provided by oceanic phytoplankton [3, 44]. However, the rate at which pelagic phytoplankton is delivered to intertidal habitats is highly variable, depending on both the concentration of offshore phytoplankton, which can vary by more than two orders of magnitude in upwelling systems, and the variability in pelagic-intertidal mixing rates, which are driven by a combination of waves, tides, and wind-driven currents. Traditional ATN modeling using a stationary approach or a constant rate of subsidy does not capture the population dynamics that result from variable phytoplankton input to the network nor the food web level consequences of the different pelagic-intertidal mixing rates. Our work advances understanding of food web responses to time-dependent variability in primary productivity by developing a novel ATN methodology that uses empirical time-series of chlorophyll-a to realistically simulate time-varying biomass of offshore phytoplankton under different pelagic-intertidal mixing scenarios to link empirical pelagic phytoplankton variability to the near-shore phytoplankton productivity that is consumed by the intertidal food web. This methodology allowed us to discover differential responses among species to the variability in phytoplankton productivity and the different pelagic-intertidal mixing rates depending on their position in the food web and relate those responses to oceanographic events captured in the empirical time-series of chlorophyll-a.

A principal, novel finding is that pulsed, time-dependent subsidy of phytoplankton can influence transient food web dynamics and lead to important temporal fluctuation in species’ biomasses. Moreover, the local rate of delivery of pelagic phytoplankton to the intertidal can amplify or alleviate the magnitude of these fluctuations, having important consequences on species’ risk of extinction as illustrated by our results with *Onchidella*. High delivery rates essentially increase the magnitude of the subsidy pulse and thus amplify the magnitude of transient effects leading to greater temporal variability in populations. Our results reinforce that of others [43] that report carnivores, as a multispecies functional group, are particularly sensitive to perturbations, showing the overall greatest cumulative amplification of variation in biomass fluctuations due to elevated phytoplankton abundances. For intermediate species, this can lead to intensification and further amplification of top-down effects. For herbivore species not buffered by increased bottom-up effects, the boost in phytoplankton pulse size can lead to sufficient intensification of top-down, negative effects that it pushes the population over a threshold of low biomass to a point that it cannot recover. For these, variation in risk to extirpation depends on network connectivity and predator identity. In contrast, under similar pelagic phytoplankton abundance, lower mixing rates can rescue these populations from extinction by favoring more balanced top-down/bottom-up effects that dampen biomass fluctuations.

Consistent with the work by Avila-Thieme *et al*. [6] using the same network model but with constant subsidy rates, we found that increased phytoplankton biomass decreases the biomass of non-planktonic producers via competition for the shared carrying capacity among producers. Additionally, both models found that increased phytoplankton biomass increased the biomass of filter feeders and that of their predators but decreased that of herbivores via the combined effects of increased predation (more abundant predators shared with filter feeders) and decreased resource availability (non-planktonic producers). Finally, both models found that omnivore species demonstrate greater resilience to elevated phytoplankton levels than herbivores by exploiting additional resources that are not affected by the decline in non-planktonic producers. This consistency between the two models suggests that these food web responses to increased phytoplankton subsidy are robust to the transient dynamics caused by the time-dependent pulsing of subsidies. Rather, these transient dynamics predict species extirpations, especially at higher trophic levels, that the model does not show using constant subsidy rates. We note these extirpations occur despite model baseline phytoplankton levels that support all species in steady state. Increased risk of extinction under transient dynamics is particularly important for understanding the effects that extreme oceanic events driven by climate change may cause to intertidal ecosystems.

The ATN model as used here only accounts for network structure, bioenergetic and trophic relationships between species, and empirical time-series data on chlorophyll-a concentration. The incorporation of other relevant biological processes such as recruitment of larval filter feeders and other invertebrates, direct competition among consumers, foreign species invasions, or habitat provisioning by basal and sessile species may improve simulations in future work. Additionally, empirical measurements that help inform the choice of pelagic-intertidal mixing rate or the proportion of ocean-derived phytoplankton in intertidal communities will significantly improve the realism of our model scenarios. Further, to fully assess the vulnerability of food webs to time-varying subsidies, there is a need to account for the different time scales of specific trophic processes in relation to the time scales of environmental subsidies. For example, time-dependent changes in feeding and assimilation processes are not included in our model. Feeding and the capacity to capitalize on accessible prey can exhibit substantial variability owing to alternative saturation rates or shifts in behavior and foraging tactics aimed at striking a balance between risk and reward in the presence of multiple stressors. Additionally, differences in the timing of resource peaks may further contribute to the intricacies of this variability. Thus, future work to incorporate the relative timing of plankton subsidy to that of temporal variability of foraging could prove insightful to the sensitivity of propagating effects and whether timing-mismatches are enough to alter the relative balance of top-down and bottom-up effects.

Our study introduces a novel methodology incorporating empirical timeseries data into the ATN framework to study the effects of variable abundances of oceanic phytoplankton on species in a Chilean rocky intertidal ecosystem. Our results draw attention to the importance of transient population dynamics and the role they may play in predicting species extirpations during extreme events. Specifically, large fluctuations in offshore phytoplankton biomass put intertidal species at risk of local extirpation even though levels of intertidal phytoplankton were not allowed to drop below a level that allowed for the persistence of all species in steady-state simulations. Our results suggest that for a species to persist in the face of large-scale biomass variability introduced by bottom-up processes, that a concomitant top-down, network-scale process is required to dampen biomass oscillations allowing for species persistence.

## Methods

### Chlorophyll-a Dataset and Preparation

Daily chlorophyll-a fluorescence measurements were taken near Las Cruces, Chile, at the Estación Costera de Investigaciones Marinas (ECIM). Raw chlorophyll-a fluorescence measurements were obtained between Jan 1999 and Dec 2010 [27, 51]. Fluorescence values were converted to biomass equivalents appropriate for the ATN framework by scaling the raw data such that the mean fluorescence over the 11-year dataset was equal to the rate of the plankton subsidy present in the original Las Cruces ATN [6]. This scaling gives the same total subsidy as the original model assuming k_mixing_ = 1.0hr^−1^ and assuming low food web phytoplankton after each timestep. We applied a nonnegative, cubic regression spline to each year to generate a continuous dataset for incorporation into the ATN framework. Years with less than 70% data coverage and years with more than 15 local extirpations (typically resulting from long periods with low recorded phytoplankton abundance) were excluded from most analyses.

### Allometric Trophic Network

The Las Cruces food web contains 106 species with 1,362 trophic links (Fig. 1A, ref. [52]). The ATN model for this network requires a constant plankton subsidy of 7,430g/m^2^ per time step [6]. ATN modeling utilizes two sets of differential equations to describe the biomass of each producer (Eq. 1) and consumer (Eq. 2) species according to the principles of mass balance as follows:

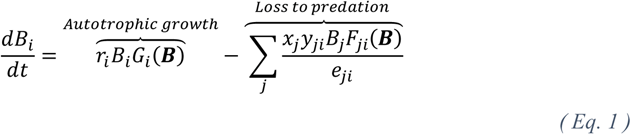

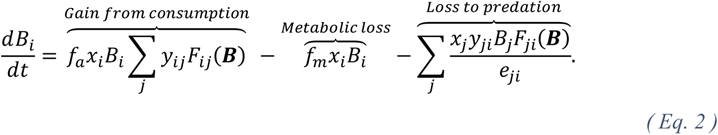

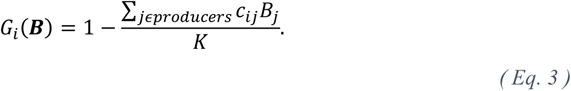

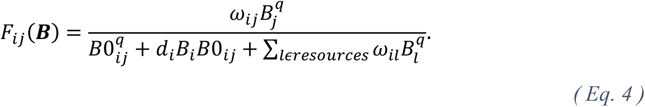

The biomass of producer species changes according to the balance of autotrophic growth with predation (Eq. 1). Autotrophic growth is controlled by the intrinsic growth rate, r_i_, and current biomass, B_i_, and is scaled by a logistic growth factor, G_i_ (Eq. 3). The logistic growth factor takes into consideration the shared, community-level, carrying capacity for producer species, K, and the interspecific competition coefficient between producers, c_ij_, to slow producer growth as the summed producer biomass approaches or exceeds carrying capacity. Biomass loss to predator j from prey i increases with the predator’s mass-specific metabolic rate, x_j_, and prey-specific attack rate, y_ji_, and decreases with the assimilation efficiency of prey j into predator i, e_ji_.

The biomass of consumer species changes according to the balance of consumer growth rate, explicitly modeled metabolic loss and mortality, and loss to predation. Consumer growth rate is modeled as assimilation of prey biomass; consequently, consumer growth rates are a function of their total consumption rate, *x_i_B_i_* ∑*_jɛprey_y_ij_F_ij_* (***B***), scaled by an assimilation coefficient, f_a_. Metabolic loss including mortality is directly proportional to current biomass and metabolic rate and is scaled by a mortality coefficient, f_m_. Finally, the loss to predation term is identical to that of producers.

The Holling’s functional response, F_ij_, is used to determine the consumption rate of each resource for each consumer (Eq. 4). This function considers the relative preference of consumer i for resource j, ω_ij_, as well as the Holling’s coefficient, q, to determine curve shape. We invoke an intermediate Holling’s Type II and Type III response by using q = 1.2 [54]. See supplementary information (Supplementary Table S1) for additional details regarding model parameterization.

All equations, parameters, and initial conditions are identical to those used in ref. [6] aside from the following modifications:

1. We created a new node titled offshore phytoplankton (see bottom node, OP, in Fig. 1A). The biomass for offshore phytoplankton, *B_op_*, is equal to the scaled, empirical dataset described above. This node connects to the food web only by water-borne exchange with the food web phytoplankton node. This introduces a new rate constant, k_mixing_, which controls the rate at which plankton moves from offshore into the food web. k_mixing_ is assigned values of 0.1, 1.0, and 10hr^−1^ as denoted throughout.
2. We separated the existing phytoplankton node into two nodes, baseline phytoplankton and food web phytoplankton. These two nodes share identical connections within the food web. Baseline phytoplankton supplies the minimal amount of phytoplankton required to prevent network collapse under steady state conditions. Baseline phytoplankton, which includes other particulate organic matter like detritus, is constant with a biomass of 3,750 g/m^2^. (All species in the network persist at this value of initial biomass. Initial biomass values of 3,700 g/m^2^ and lower cause multiple species extinctions within only a handful of timesteps.) The food web phytoplankton node provides the primary supply of phytoplankton to consumers in the food web. In addition to the ATN modeling of producer species listed in Eq. 1, the biomass of food web phytoplankton, *B_fp_*, is subsidized by biomass from offshore phytoplankton via water-borne exchange (Eq. 5).

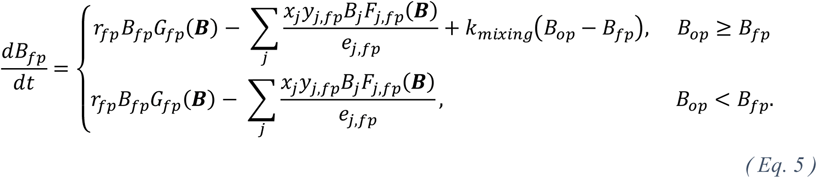

### Simulations

Each year of empirical data, from Jan 1 to Dec 31, was analyzed independently. To better compare simulations performed under different empirical datasets, we sought to reduce the transient dynamics introduced by having initial conditions that were significantly different from runtime conditions. We evaluated two separate strategies for reducing the impacts of initial conditions and found little change between the two. Strategy 1 involved running simulations under an “equilibration phase” with the offshore phytoplankton node deactivated for ten years of simulation time prior to activating the empirical data during the “experimental phase” (Fig. S4). Strategy 2 replaces the “equilibration phase” with ten appended copies of the empirical dataset. This allows population biomasses to more gently approach runtime conditions than strategy 1. Population distributions were highly similar between each strategy, and the results presented here were obtained using strategy 1.

### Coefficient of Variation and Violin Plots

We use the coefficient of variation, the ratio of standard deviation to mean, as the preferred metric of variability for the timeseries data presented here. We calculated coefficient of variation for each species during the time interval when its biomass is above the extinction threshold of 10^−6^g/m^2^ (i.e., trailing zeroes were truncated where applicable).

Violin plots were used to display distributional data on coefficients of variation. These plots are a modification of box plots in that they show the median value, each individual value, interquartile range, and shade the kernel density estimate to give an indication of the data distribution. Wider sections of violin plots represent a higher probability that members in the population will take on a given value. Violin plots were generated in Matlab using code from Bastian Bechtold [53].

## Acknowledgements

We thank Kayla R. S. Hale, M. Isidora Ávila-Thieme, Sophia M. Simon, and Sabine Dritz for insightful discussions pertaining to programming, model parameterization, and interpretation of results.

This study was supported by National Science Foundation grant DEB-2224915 and UC Davis Seed Grant for Advancing Sustainable Development Goals.

## Author Contributions

C.S.D., J.L.L. and F.S.V. conceived the study. E.A.W. compiled empirical datasets. C.S.D. cleaned datasets, modified the dynamic model, implemented simulations, and prepared figures. C.S.D., J.L.L., and F.S.V. analyzed simulation results. C.S.D. and F.S.V. wrote the first draft of the manuscript. All authors conducted revisions and contributed to the final version of the paper.

## Data availability statement (mandatory)

Simulation code and the Chilean intertidal data will be available upon acceptance at the repository https://github.com/Valdovinos-Lab/Chilean_Variable_Subsidy. The Chilean intertidal food web and species body sizes can also be found in 12.

## References

1. Krenz, C. et al. Ecological subsidies to rocky intertidal communities: Linear or non-linear changes along a consistent geographic upwelling transition?. Journal of Experimental Marine Biology and Ecology 409, 361–370 (2011). https://doi.org/10.1016/j.jembe.2011.10.003.

2. Leslie, H. M., et al. Barnacle reproductive hotspots linked to nearshore ocean conditions. Proceedings of the National Academy of Sciences 102, 10534–10539 (2005). https://doi.org/10.1073/pnas.0503874102.

3. Menge, B. A. et al. Coastal oceanography sets the pace of rocky intertidal community dynamics. Proceedings of the National Academy of Sciences 100, 12229–12234 (2003). https://doi.org/10.1073/pnas.1534875100.

4. Phillips, N. E. Growth of filter-feeding benthic invertebrates from a region with variable upwelling intensity. Marine Ecology Progress Series 295, 79–89 (2005). https://doi.org/10.3354/meps295079.

5. Phillips, N. E. A spatial gradient in the potential reproductive output of the sea mussel Mytilus californianus. Marine Biology 151, 1543–1550 (2007). https://doi.org/10.1007/s00227-006-0592-x.

6. Ávila-Thieme, M. I. et al. Alteration of coastal productivity and artisanal fisheries interact to affect a marine food web. Scientific Reports 11, 1–14 (2021). https://doi.org/10.1038/s41598-021-81392-4.

7. Bracken, M. E. et al. Mussel selectivity for high-quality food drives carbon inputs into open-coast intertidal ecosystems. Marine Ecology Progress Series 459, 53–62 (2012). https://doi.org/10.3354/meps09764.

8. Morgan, S. G., Shanks, A. L., MacMahan, J. H., Reniers, A. J. & Feddersen, F. Planktonic Subsidies to Surf-Zone and Intertidal Communities. Annual Review of Marine Science 10, 345–369 (2018). https://doi.org/10.1146/annurev-marine-010816-060514.

9. Largier, J. L et al. WEST: A northern California study of the role of wind-driven transport in the productivity of coastal plankton communities. Deep Sea Research Part II: Topical Studies in Oceanography 53, 2833–2849 (2006). https://doi.org/10.1016/j.dsr2.2006.08.018.

10. Largier, J. L. Upwelling bays: how coastal upwelling controls circulation, habitat, and productivity in bays. Annual Review of Marine Science 12, 415–447 (2020). https://doi.org/10.1146/annurev-marine-010419-011020.

11. Largier, J. L. Rip Currents and the Influence of Morphology on Wave-Driven Cross-Shore Circulation. Reference Module in Earth Systems and Environmental Sciences, 100–121 (2022).

12. Morgan, S. G. et al. Surfzone hydrodynamics as a key determinant of spatial variation in rocky intertidal communities. Proceedings of the Royal Society B: Biological Sciences 283, 20161017 (2016). https://doi.org/10.1098/rspb.2016.1017.

13. Shanks, A. L., Morgan, S. G., MacMahan, J. & Reniers, A. J. Surf zone physical and morphological regime as determinants of temporal and spatial variation in larval recruitment. Journal of Experimental Marine Biology and Ecology 392, 140–150 (2010). https://doi.org/10.1016/j.jembe.2010.04.018.

14. Hastings, A., et al. Transient phenomena in ecology. Science 361, eaat6412 (2018). https://doi.org/10.1126/science.aat641212.

15. Blanchette, C. A., Wieters, E. A., Broitman, B. R., Kinlan, B. P. & Schiel, D. R. Trophic structure and diversity in rocky intertidal upwelling ecosystems: A comparison of community patterns across California, Chile, South Africa and New Zealand. Progress in Oceanography 83, 107–116 (2009). https://doi.org/10.1016/j.pocean.2009.07.038.

16. Gutiérrez, D., Akester, M. & Naranjo, L. Productivity and sustainable management of the Humboldt Current large marine ecosystem under climate change. Environmental Development 17, 126–144 (2016). https://doi.org/10.1016/j.envdev.2015.11.004.

17. Montecino, V. & Lange, C.B. The Humboldt Current System: Ecosystem components and processes, fisheries, and sediment studies. Progress in Oceanography 83, 65–79 (2009). https://doi.org/10.1016/j.pocean.2009.07.041.

18. Ochoa, N., Taylor, M.H., Purca, S. & Ramos, E. Intra- and interannual variability of nearshore phytoplankton biovolume and community changes in the northern Humboldt Current system. Journal of plankton research 32, 843–855 (2010). https://doi.org/10.1093/plankt/fbq022.

19. Chavez, F.P. & Messié, M. A comparison of eastern boundary upwelling ecosystems. Progress in Oceanography 83, 80–96 (2009). https://doi.org/10.1016/j.pocean.2009.07.032.

20. Daneri, G., et al. Primary production and community respiration in the Humboldt Current System off Chile and associated oceanic areas. Marine Ecology Progress Series 197, 41–49 (2000). https://doi.org/10.3354/meps197041.

21. Ryther, J.H. Photosynthesis and Fish Production in the Sea: The production of organic matter and its conversion to higher forms of life vary throughout the world ocean. Science 166, 72–76 (1969). https://doi.org/10.1126/science.166.3901.72.

22. Oyarzún, D. & Brierley, C.M. The future of coastal upwelling in the Humboldt current from model projections. Climate dynamics 52, 599–615 (2019). https://doi.org/10.1007/s00382-018-4158-7.

23. Martinez, E., Antoine, D., D’Ortenzio, F. & Gentili, B. Climate-driven basin-scale decadal oscillations of oceanic phytoplankton. Science 326, 1253–1256 (2009). https://doi.org/10.1126/science.1177012.

24. Thatje, S., Heilmayer, O. & Laudien, J. Climate variability and El Niño Southern Oscillation: implications for natural coastal resources and management. Helgoland Marine Research 62, 5–14 (2008). https://doi.org/10.1016/j.csr.2013.11.015.

25. Bakun, A. Global climate change and intensification of coastal ocean upwelling. Science 247, 198–201 (1990). https://doi.org/10.1126/science.247.4939.198.

26. Sydeman, W.J., et al. Climate change and wind intensification in coastal upwelling ecosystems. Science 345, 77–80 (2014). https://doi.org/10.1126/science.1251635.

27. Weidberg, N., et al. Spatial shifts in productivity of the coastal ocean over the past two decades induced by migration of the Pacific Anticyclone and Ba’un’s effect in the Humboldt Upwelling Ecosystem. Global and Planetary Change 193, 103259 (2020). https://doi.org/10.1016/j.gloplacha.2020.103259.

28. Yodzis, P. & Innes, S., 1992. Body size and consumer-resource dynamics. The American Naturalist 139, 1151–1175 (1992). https://doi.org/10.1086/285380.

29. Brose, U., et al. Consumer–resource body-size relationships in natural food webs. Ecology 87, 2411–2417 (2006). https://doi.org/10.1890/0012-9658(2006)87[2411:CBRINF]2.0.CO;2.

30. Berlow, E.L., et al. Simple prediction of interaction strengths in complex food webs. Proceedings of the National Academy of Sciences 106, 187–191 (2009). https://doi.org/10.1073/pnas.0806823106.

31. Boit, A., Martinez, N.D., Williams, R.J. & Gaedke, U. Mechanistic theory and modelling of complex food-web dynamics in Lake Constance. Ecology letters 15, 594–602 (2012). https://onlinelibrary.wiley.com/doi/10.1111/j.1461-0248.2012.01777.x.

32. Brose, U., Williams, R.J. & Martinez, N.D. Allometric scaling enhances stability in complex food webs. Ecology letters 9, 1228–1236 (2006). https://doi.org/10.1111/j.1461-0248.2006.00978.x.

33. Schneider, F.D., Brose, U., Rall, B.C. & Guill, C. Animal diversity and ecosystem functioning in dynamic food webs. Nature Communications 7, 1–8 (2016). https://doi.org/10.1038/ncomms12718.

34. Albert, G., Gauzens, B., Loreau, M., Wang, S. & Brose, U. The hidden role of multi-trophic interactions in driving diversity–productivity relationships. Ecology Letters 25, 405–415 (2022). https://doi.org/10.1111/ele.13935.

35. Gauzens, B., Rall, B.C., Mendonça, V., Vinagre, C. & Brose, U. Biodiversity of intertidal food webs in response to warming across latitudes. Nature Climate Change 10, 264–269 (2020). https://doi.org/10.1038/s41558-020-0698-z.

36. Glaum, P., Cocco, V. & Valdovinos, F.S. Integrating economic dynamics into ecological networks: The case of fishery sustainability. Science Advances 6, p.eaaz4891 (2020). https://doi.org/10.1126/sciadv.aaz4891.

37. Perälä, T. & Kuparinen, A. Eco-evolutionary dynamics driven by fishing: From single species models to dynamic evolution within complex food webs. Evolutionary Applications 13, 2507–2520 (2020). https://doi.org/10.1111/eva.13058.

38. Uusi-Heikkilä, S., Perälä, T. & Kuparinen, A. Fishing triggers trophic cascade in terms of variation, not abundance, in an allometric trophic network model. Canadian Journal of Fisheries and Aquatic Sciences 99, 1–11 (2022). https://doi.org/10.1139/cjfas-2021-0146.

39. Brose, U., et al. Predator traits determine food-web architecture across ecosystems. Nature Ecology & Evolution 3, 919–927 (2019). https://doi.org/10.1038/s41559-019-0899-x.

40. Petchey, O.L., Beckerman, A.P., Riede, J.O. & Warren, P.H. Size, foraging, and food web structure. Proceedings of the National Academy of Sciences 105, 4191–4196 (2008). https://doi.org/10.1073/pnas.0710672105.

41. Peters, R.H. Size structure of the plankton community along the trophic gradient of Lake Memphremagog. Canadian Journal of Fisheries and Aquatic Sciences 40, 1770–1778 (1983). https://doi.org/10.1139/f83-206.

42. Shurin, J.B., Gruner, D.S. & Hillebrand, H. All wet or dried up? Real differences between aquatic and terrestrial food webs. Proceedings of the Royal Society B: Biological Sciences 273, 1–9 (2006). https://doi.org/10.1098/rspb.2005.3377.

43. Berg, S., Pimenov, A., Palmer, C., Emmerson, M. & Jonsson, T. Ecological communities are vulnerable to realistic extinction sequences. Oikos 124, 486–496 (2015). https://doi.org/10.1111/oik.01279.

44. Menge, B.A. Top-Down and bottom-up community regulation in marine rocky intertidal habitats. Journal of Experimental Marine Biology and Ecology 250, 257–289 (2000). https://doi.org/10.1016/s0022-0981(00)00200-8.

45. Melet, A.V., Hallberg, R. & Marshall, D.P. The role of ocean mixing in the climate system. Ocean Mixing, 5–34 (2022). https://doi.org/10.1016/B978-0-12-821512-8.00009-8.

46. Xiu, P., Chai, F., Curchitser, E.N. & Castruccio, F.S. Future changes in coastal upwelling ecosystems with global warming: The case of the California Current System. Scientific Reports 8, 1–9 (2018). https://doi.org/10.1038/s41598-018-21247-7.

47. Kaplan, D.M., Largier, J. & Botsford, L.W. HF radar observations of surface circulation off Bodega Bay (northern California, USA). Journal of Geophysical Research: Oceans 110, C10 (2005). https://doi.org/10.1029/2005JC002959.

48. Cavole, L.M., et al. Biological impacts of the 2013–2015 warm-water anomaly in the Northeast Pacific: winners, losers, and the future. Oceanography 29, 273–285 (2016). https://doi.org/10.5670/oceanog.2016.32.

49. Menge, B. A., et al. Climatic variation alters supply-side ecology: impact of climate patterns on phytoplankton and mussel recruitment. Ecological Monographs 79, 379–395 (2009).

50. Navarrete, S. A., Barahona, M., Weidberg, N. & Broitman, B. R. Climate change in the coastal ocean: shifts in pelagic productivity and regionally diverging dynamics of coastal ecosystems. Proceedings of the Royal Society B: Biological Sciences 289, 20212772 (2022).

51. Wieters, E.A., et al. Alongshore and temporal variability in chlorophyll a concentration in Chilean nearshore waters. Marine Ecology Progress Series 249, 93–105 (2003). https://doi.org/10.3354/meps249093.

52. Kéfi, S., et al. Network structure beyond food webs: mapping non-trophic and trophic interactions on Chilean rocky shores. Ecology 96, 291–303 (2015).

53. Bechtold, B., Violinplot-Matlab, (2022), github, https://github.com/bastibe/Violinplot-Matlab.

54. Williams, R. J. Effects of network and dynamical model structure on species persistence in large model food webs. Theor. Ecol. 1, 141–151 (2008).

