## Supplementary Information for "Modeling time-varying phytoplankton subsidy reveals at-risk species in a Chilean intertidal ecosystem"

### Supplemental Materials

**Supplementary Table S1** – Table describing empirical datasets and simulation results performed under $k_{mixing} = 1.0 hr^{-1}$ (intermediate pelagic-intertidal mixing rate). Table shows year, number of days with data collection, percentage of year with data coverage, largest gap in data collection measured in days, the number of species predicted to have gone locally extinct, and their network ID. Supplementary Table S2 details these species.

| Year | #days with data | %coverage | Biggest gap | Number Extinct with $k_{mix} = 1$ | Identities of extinct spec |
|---|---|---|---|---|---|
| 1999 | 270 | 79.97 | 14 | 6 | 19, 29, 98, 99, 100, 106 |
| 2000 | 201 | 55.07 | 5 | 10 | 2, 3, 19, 28, 29, 96, 98, 99, 100, 106 |
| 2001 | 242 | 66.3 | 40 (start) | 6 | 2, 3, 19, 28, 96, 106 |
| 2002 | 278 | 76.16 | 4 | 10 | 2, 3, 19, 28, 29, 96, 98, 99, 100, 106 |
| 2003 | 289 | 79.18 | 4 | 6 | 19, 29, 98, 99, 100, 106 |
| 2004 | 280 | 76.71 | 7 (start) | 6 | 2, 3, 19, 28, 96, 106 |
| 2005 | 285 | 78.08 | 6 | 2 | 19, 106 |
| 2006 | 248 | 67.95 | 32 (start) | 9 | 2, 3, 19, 28, 96, 98, 99, 100, 106 |
| 2007 | 251 | 68.77 | 5 | 10 | 2, 3, 19, 28, 29, 96, 98, 99, 100, 106 |
| 2008 | 241 | 66.03 | 35 (start) | 5 | 2, 3, 19, 28, 96 |
| 2009 | 250 | 68.49 | 6 | 5 | 2, 3, 19, 28, 96 |
| 2010 | 229 | 62.74 | 13 | 5 | 2, 3, 19, 28, 96 |

**Supplementary Table S2** – Table describing species that predicted to have gone extinct in simulations performed under $k_{mixing} = 1.0hr^{-1}$. For each species, the table lists network ID, scientific name, colloquial name, trophic guild, and number of years in which it was simulated to have gone extinct.

| ID | Name | Colloquial Name | Guild | #years extinct with $k_{mix} = 1$ |
|---|---|---|---|---|
| 2 | acanthocyclus gayi | crab | carnivore | 9 |
| 3 | acanthocyclus hassleri | crab | carnivore | 9 |
| 19 | heliaster helianthus | sun star | carnivore | 12 |
| 28 | stichaster striatus | starfish | carnivore | 9 |
| 29 | tegula atra | sea snail | herbivore | 5 |
| 96 | gulls | bird | carnivore | 9 |
| 98 | echinolittorina peruviana | sea snail | herbivore | 6 |
| 99 | austrolittorina araucana | sea snail | herbivore | 6 |
| 100 | onchidella sp | sea snail | herbivore | 6 |
| 106 | trimusculus peruvianus | sea snail | herbivore | 9 |

**Supplementary Table S3 -** Initial parameter values and equilibrium parameter values (see Fig S4) used in our version of the Allometric Trophic Network (ATN) model of the central Chilean coast. All initial parameter values are taken from Ávila-Thieme 2021.

| Parameter | Units | Definition | Initial and Equilibrium Values min, max |
|---|---|---|---|
| B | $g / m^2$ | Population density | IV: $1.24 \times 10^{-4}$, 112,107<br>EV: $3.26 \times 10^{-4}$, 4,401.6 |
| r | $day^{-1}$ | Mass-specific growth rate of producers | 0.1075, 3.76 |
| x | $day^{-1}$ | Mass-specific growth rate of consumers | 0.7284, 70.96 |
| G | - | Logistic growth factor | IV: 0, 0.2495<br>EV: $8.99 \times 10^{-4}$, 0.0259 |
| y | - | Maximum consumption rate | 1, 5.8 |
| F | - | Functional response | IV:$1.23 \times 10^{-11}$, 0.7154<br>EV: $1.17 \times 10^{-8}$, 0.3622 |
| $f_a$ | - | Fraction of biomass assimilated by consumers | 0.4 |
| $f_m$ | - | Fraction of biomass lost to metabolic maintenance | 0.1 |
| e | - | Assimilation efficiency | 0.45, 0.85 |

**Supplementary Table S4** – Coefficients of variation were calculated for summed biomass across each trophic grouping above for the full year datasets of 1999-2010, and for each of the three values of $k_{mixing}$. Coefficients were normalized to offshore phytoplankton. Reported values give average and standard deviation over the 12 years.

| | | $k_{mixing}$ ($hr^{-1}$) | | |
|---|---|---|---|---|
| | | 0.1 | 1.0 | 10 |
| Trophic Group | Offshore Phytoplankton | 1.00 ± 0.00 | 1.00 ± 0.00 | 1.00 ± 0.00 |
| | Foodweb Phytoplankton | 0.20 ± 0.08 | 0.39 ± 0.12 | 0.54 ± 0.17 |
| | Non-Planktonic Producers | 0.01 ± 0.00 | 0.02 ± 0.02 | 0.04 ± 0.04 |
| | Filter Feeders | 0.21 ± 0.11 | 0.46 ± 0.16 | 0.66 ± 0.23 |
| | Herbivores | 0.07 ± 0.03 | 0.14 ± 0.07 | 0.16 ± 0.08 |
| | Omnivores | 0.01 ± 0.00 | 0.03 ± 0.02 | 0.13 ± 0.15 |
| | Consumers | 0.52 ± 0.23 | 0.78 ± 0.13 | 1.06 ± 0.17 |

**Supplementary Figure S1** – Results shown mirror those in Figure 2. Each year has four associated panels – "Phytoplankton Abundance" shows biomass of both offshore phytoplankton (OP) and Food web Phytoplankton (FP); "Barnacle Abundance" shows biomass of the barnacle *B. laevis* (BL); "Producer Abundance" shows the summed biomass of all producer species (Prod) and the summed biomass of all non-planktonic producer species (NP Prod); and "Sea Snail Abundance" shows biomass of the omnivore sea snail *L. orbignyi* (LO) and the herbivore sea snail *T. atra* (TA). All panels show results using an intermediate pelagic-intertidal mixing rate parameter, $k_{mixing} = 1.0hr^{-1}$.

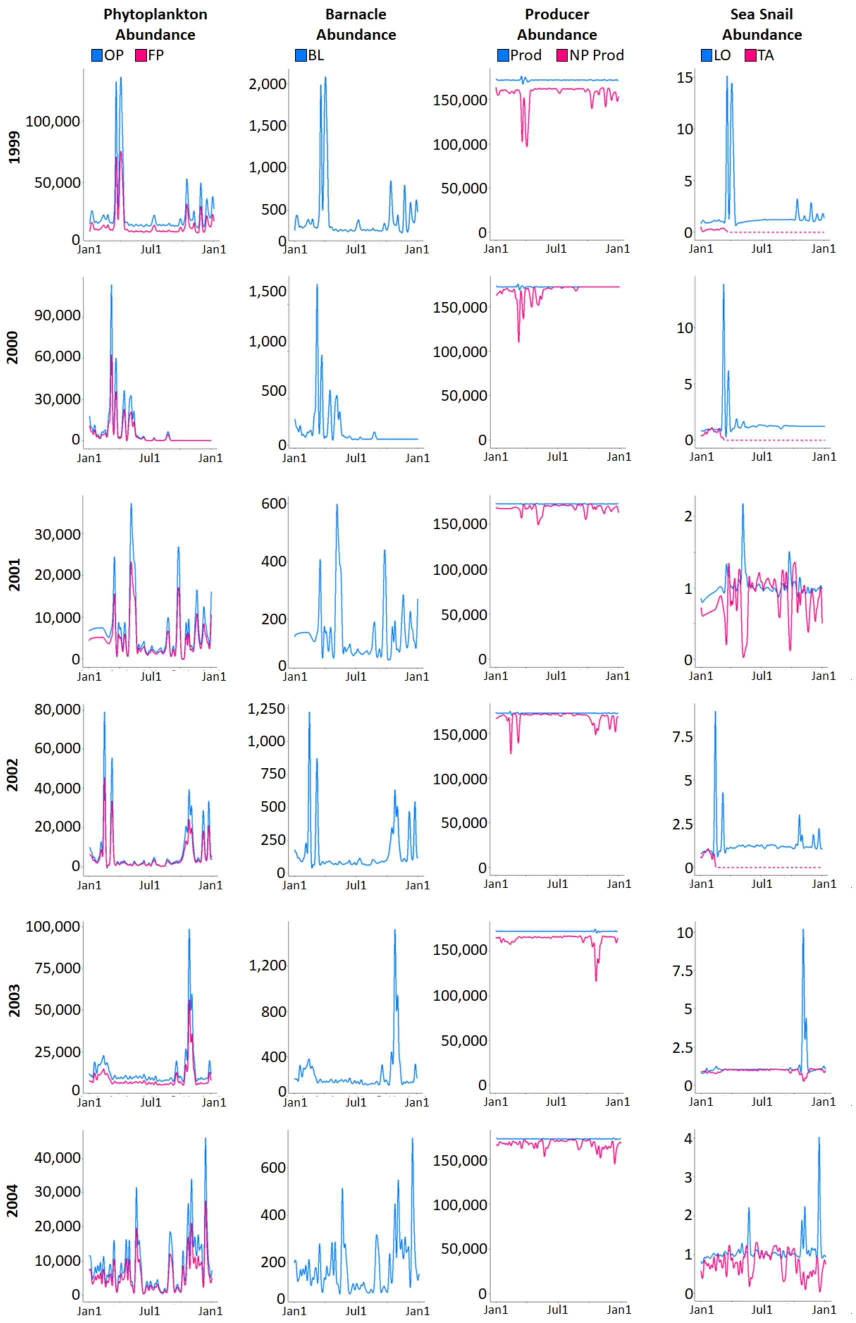

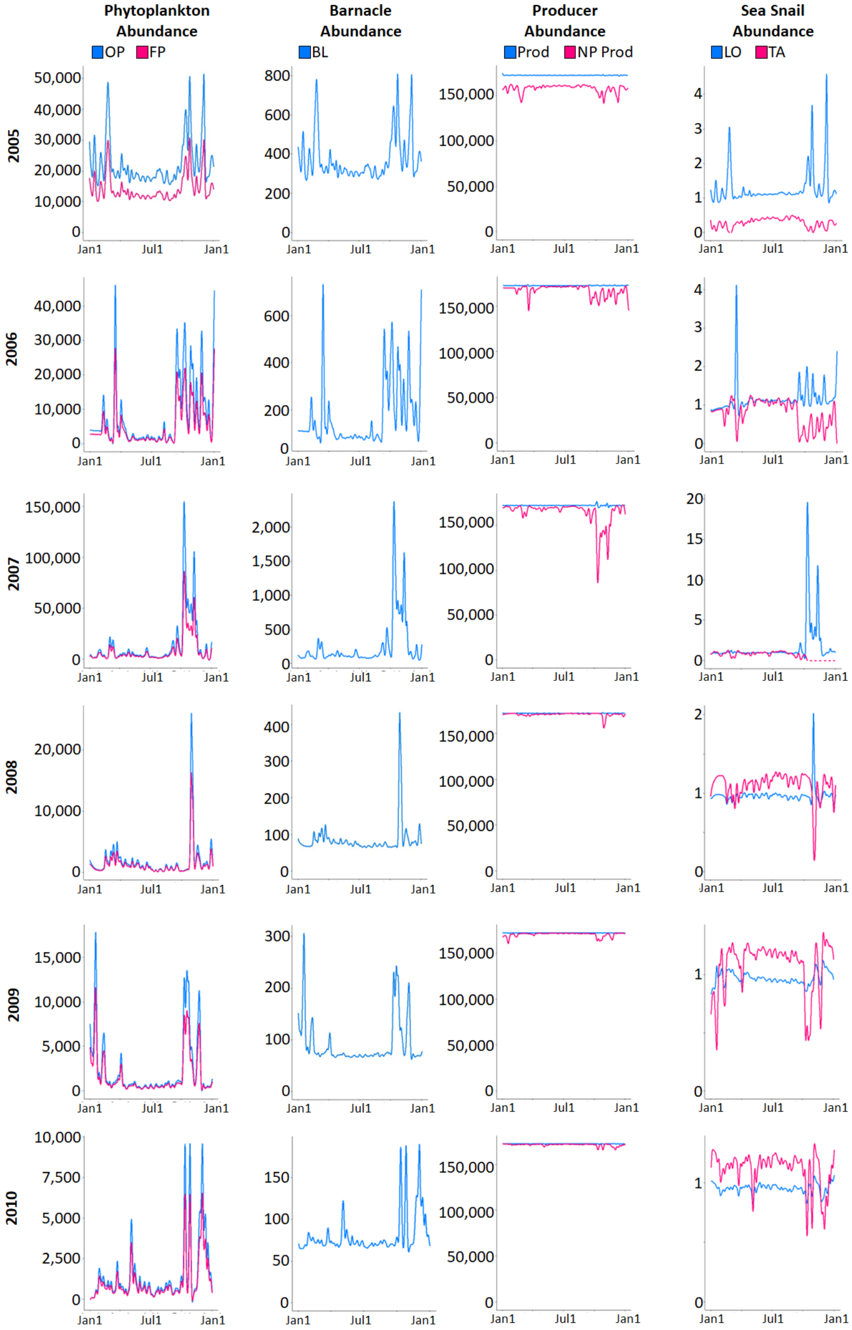
Phytoplankton Abundance
OP
FP
Barnacle Abundance
BL
Producer Abundance
Prod
NP Prod
Sea Snail Abundance
LO
TA
2005
2006
2007
2008
2009
2010
Jan1
Jul1
Jan1

**Supplementary Figure S2** – Relative abundance of all 15 filter feeders as they responded to offshore phytoplankton in 2003 presented in Figure 2. Each species' biomass was normalized to its baseline level. These curves include 5 barnacles, 3 mussels, 1 worm, 1 tunicate, and 5 crabs. All biomass curves show positive responses to elevated offshore phytoplankton.

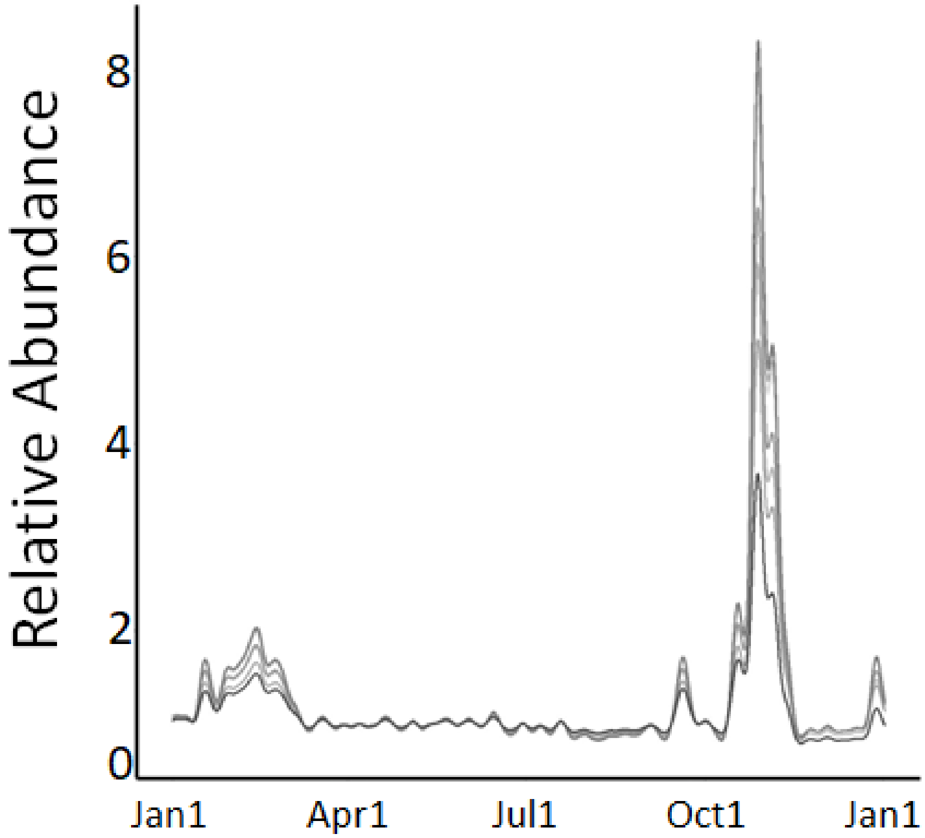

**Supplementary Figure S3** – Relative abundance of all 20 sea snails as they responded to offshore phytoplankton in 2003 presented in figure 2. Each species' biomass was normalized to its baseline level. These curves include 16 omnivorous sea snails (blue curves) and 4 herbivorous sea snails (green curves). All omnivorous sea snails show positive responses to elevated offshore phytoplankton, while all herbivorous sea snails show negative responses to elevated offshore phytoplankton.

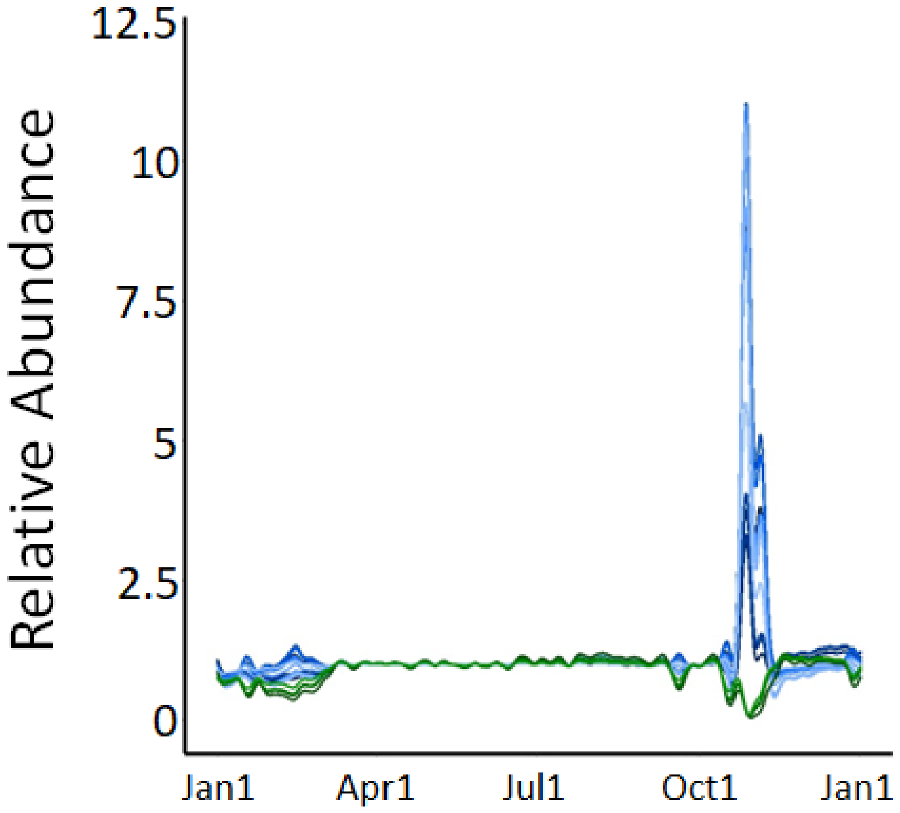

**Supplementary Figure S4** – Results shown mirror those in Fig. 4AB and are summarized in Table 1. Each year has two associated figures – top panel: offshore phytoplankton biomass, bottom panel: simulated *Onchidella* biomass under $k_{mixing} = 0.1 hr^{-1}$ (teal), $1.0 hr^{-1}$ (green), and $10 hr^{-1}$ (navy). Time-axes were truncated to show 3 month periods, with at least 1 month on either side of extirpation events. 2009 and 2010 show results from the full year because neither year showed extirpations.

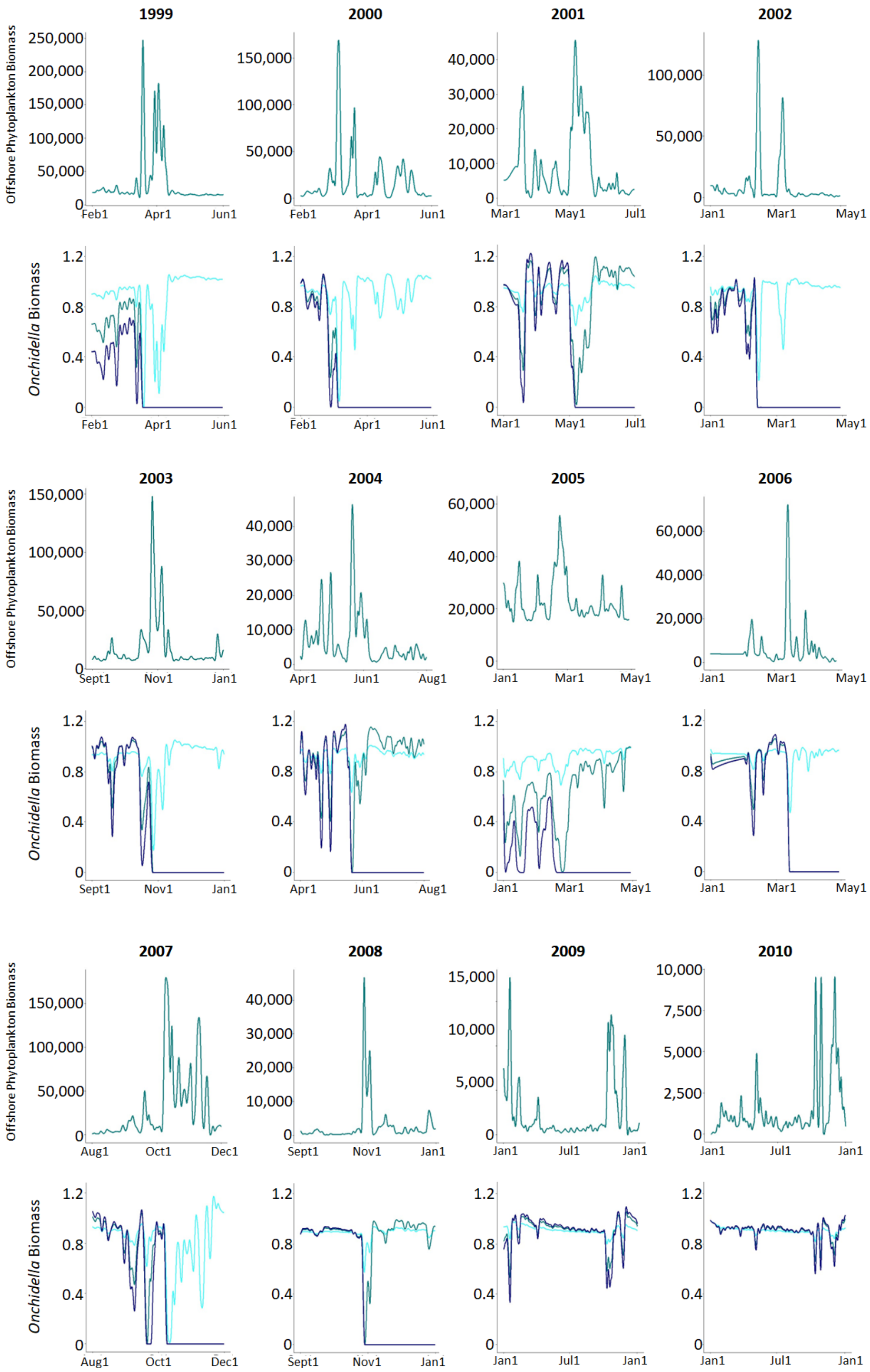

**Supplementary Figure S5** – Simulation strategy highlighting ‘equilibration period’ followed by ‘treatment period’. (A) Baseline phytoplankton is active for one year of simulation time with food web phytoplankton and offshore phytoplankton deactivated. Offshore phytoplankton is then activated and one year of empirical offshore phytoplankton data is simulated. (B) The resulting biomass curves of all 105 non-phytoplankton species.

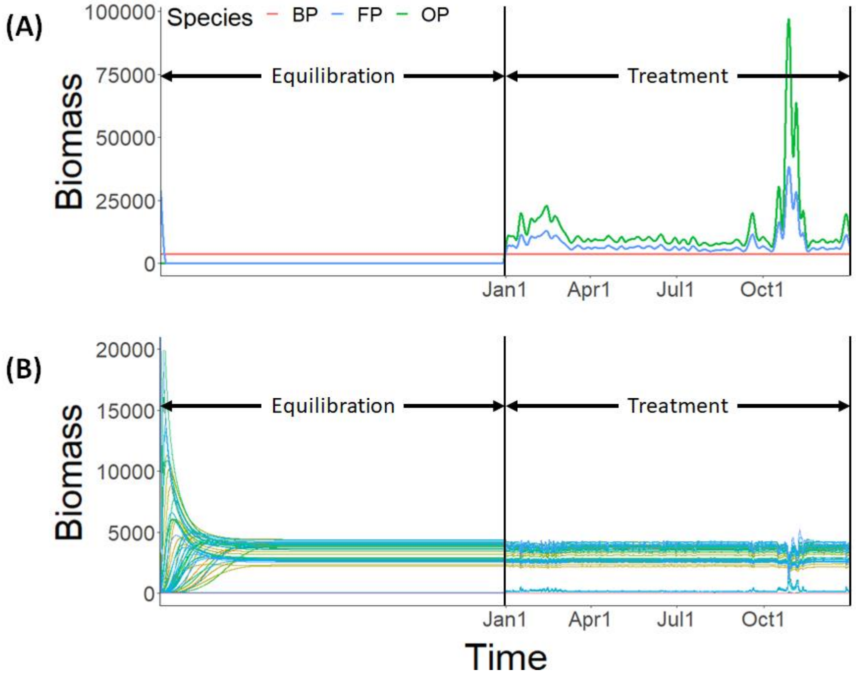
